# The kinematics and aerodynamics of hinged wing of honeybees during takeoff

**DOI:** 10.1101/2024.01.30.577934

**Authors:** Yizhe Li, Zhanzhou Hao, Biao Yang, Weijie Gong, Bo Yin, Chao Wang, Jialei Song, Ling Yin

**Affiliations:** School of Mechanical Engineering, Dongguan University of Technology, Dongguan, Guangdong, China; College of Mechatronics and Control Engineering, Shenzhen University, Shenzhen, Guangdong, China; School of Engineering Science, University of Chinese Academy of Sciences, Beijing, China; Key Laboratory of Mechanics in Fluid Solid Coupling Systems, Institute of Mechanics, Chinese Academy of Sciences, Beijing, China; Guangdong Provincial Key Laboratory of Intelligent Disaster Prevention and Emergency Technology, Dongguan University of Technology, Dongguan, Guangdong, China

**Keywords:** hinged wing, honeybee takeoff, chordwise deformation, flapping actuation

## Abstract

In this study, we utilize a high-speed camera array to meticulously capture the intricate wing kinematics of honeybees during free voluntary takeoffs. This allows us to investigate the variation in kinematic parameters over time. According to the variation of Euler angles, the takeoff process can be categorized into three stages: initial, adjusting, and stable. Our analysis reveals that honeybees typically execute at least 15 wingbeats before taking off, with wing stroke amplitudes exceeding 100 degrees and wingbeat frequencies ranging from 180 to 260 Hz. Significantly, the hinged wing mechanism, connecting the forewing and hindwing via hamuli, undergoes considerable chordwise deformation during this process, with the angle between the fore- and hind-wings surpassing 50 degrees. Notably, the forewing and hindwing maintain a positive camber throughout the wingbeat cycle during takeoff, contributing to the enhanced thrust generation instead of lift, comparing to the flat wings. The positive camber can be passively formed at beginning and ending of each half stroke, but should be actively maintained around middle half-stroke.This study provides valuable insights for aeronautical engineers in the design of flapping wing micro air vehicles, specifically in the effective implementation of hinged wings inspired by the wing motion of honeybees.

## 1 Introduction

Takeoff, as the first step launching to the air, is a complex process that plays an crucial role in escaping predators, foraging and finding mates for flying insects^[1, 2]^. When these insects take off, they generate an upward force to counteract gravity using either legs or wings. According to the apparatus that used for takeoff, insect takeoff can be categorized into three types: leg-assisted (or jumping) takeoff, wing-assisted takeoff and the combined takeoff^[3-6]^. Different insects use varying methods of takeoffs, the same insect might also use different ones depending on the situation. For instance, fruit flies exhibit at least two distinct types of takeoffs: leg-assisted escape takeoff, triggered by visual stimuli, and voluntary takeoff. In the case of escape takeoff, the fruit fly applies a much greater force with its legs, enabling faster takeoff, whereas voluntary takeoff results in slower and more stable flight^[7]^. Wingless or leg-assisted insects like fleas, locusts or hoppers use their powerful legs to jump rapidly into the air, which is usually featured with a short duration, high acceleration, emergence-involvement and uncontrollable trajectory^[4, 8-10]^. Meanwhile, the wing-assisted takeoff usually features volunteer takeoff, ground effect and controllable trajectory^[11, 12]^.

In winged insects, the involvement of legs and wings occurs so quickly that it is difficult to distinguish with the naked eyes. Using a high speed camera to study grasshopper takeoff, researchers observed that the grasshopper’s body has already lifted several body lengths above the takeoff platform before the wings begin their flapping motion, with an average time delay for wing flapping of 33.2±12.8 ms^[13]^. Analysis of butterflies revealed that a leg impulse creates upward motion during takeoff, with the horizontal and vertical vortex rings being generated for aerodynamic lift and thrust during both downstroke and upstroke^[14]^. Further studies found that takeoff is primarily driven by substantial aerodynamic forces generated in the first half of the downstroke of the wings, with the “clap and fling” mechanism -a form of interference effect between initially narrowly separated wings-being the main cause of these significant aerodynamic forces^[15]^. During takeoff of a dronefly, the aerodynamic force steadily increases from zero to slightly greater than the dronefly’s body weight, while leg force monotonically decreases from being equal to the dronefly’s body weight, while standing, to zero. This indicates that droneflies achieve takeoff without any jumping behavior, relying solely on wing flapping^[3]^. With numerical simulation, Bode-Oke discovered a damselfly generates about three times its body weight during the first half-stroke for liftoff, as a leading edge vortex (LEV) is formed during both the downstroke and upstroke on all the four wings^[16]^. However, the LEV is reduced in the subsequent upstrokes following takeoff, leading to a drastic reduction in the magnitude of the aerodynamic force.

The flow around the flapping wing of insects is unsteady, and the aerodynamic force primarily comes from the unsteady mechanisms, such as “delayed stall” ^[17]^, wake capture^[18]^, and “clap and fling” ^[19]^. The flapping wing, as the low Reynolds number and unsteady flow being around it, leads to some high lift production mechanisms. The study on mosquitoes revealed that the small amplitude of high-frequency flap of mosquitoes did not have a dominant effect on the generation of lift due to the leading edge vortex, while the drag generated by the trailing edge vortex and rotation process also produced significant weight support ^[20]^. A recent study on tiny beetles with bristly wings generate significant resistance to lift and fly through the movement of a horizontal 8 shape^[21]^. As the takeoff of insects usually requires the lift much larger than the weight, the wing-assisted takeoff might exist some new mechanisms of high lift generation.

Previous studies primarily focus on the unsteady aerodynamics of flat wings while the deformation based on real elasticity is scarce. Due to the distribution of wing veins and thickness changes of folds, the deformation of insect wings during flapping is mainly torsion in the spanwise direction and curvature in the chord direction^[22, 23]^. The stiffness of the anisotropy wing in the spanwise direction is 1-2 orders of magnitude greater than that in the chord direction^[24]^. However, in some four-winged insects, the fore- and hindwings are coupled during flight, with the leading edge of the hindwing are ventrally locked with the retinaculum at the trailing edge of the forewing for moths^[25]^, or the costal margin of the hindwing base and overlaps with the trailing edge of the forewing base for butterflies^[26]^. As a results, the chord bending in this type of forewing and hindwing coupling is significant. Ma et al. used a high-speed camera to capture the hovering flight of a bee, observing the angle changes between the forewing and hindwing during the flapping ^[27]^.

It was found that the angle changes of the flexible model led to an increase in lift and drag, and the rate of drag increase was higher. However, the kinematics reconstructed was two-dimensional, and the new high lift mechanisms potentially due to the three-dimensional effect might be neglected. A three dimensional study on honeybee flight showed its high frequency and low stroke amplitude flapping wings generate prominent force peaks during the beginning, middle and end of each stroke, indicating the importance of additional unsteady mechanisms^[28]^. However, the wing model used is flat, overlooking the significant chordwise deformation due to forewing and hindwing interconnection. Therefore, in order to further deepen the understanding of the mechanism of bee takeoff and forewing and hindwing coupling motion, this study used four high-speed cameras to capture the three-dimensional kinematics of takeoff of honeybees, and analyzed the takeoff process and forewing and hindwing coupling from both kinematic and aerodynamic perspectives.

## 2 Methods

### 2.1 Subjects

The honeybees, *Apis cerana*, used in the experiment are captured from a beekeeping facility near the laboratory and the observations are done within four hours of the capture. After completing the flight video collection of the bees, their mass is measured using an electronic analytical balance (AP135W, *Shimadzu*, Japan, with an accuracy of 0.01 mg). The average mass of honeybees is 69.63 ± 1.66 mg (mean ± standard error of the mean, s.e.m.). The geometric dimensions of the wings have also been measured and listed in Table 1. In this study, the takeoff of the honeybees is voluntary, with no external stimulus throughout the measurement.

**Table 1.** Static parameters of the honeybee, *Apis cerana*.

| Parameters | Values |
| --- | --- |
| Mass (mg) | 69.63±1.66 |
| Forewing area (mm<sup>2</sup>) | 16.19±0.06 |
| Hindwing area (mm<sup>2</sup>) | 8.31±0.11 |
| Forewing length (mm) | 8.78±0.07 |
| Hindwing length (mm) | 6.09±0.07 |
| Forewing mean chord length (mm) | 1.84±0.01 |
| Hindwing mean chord length (mm) | 1.36±0.02 |

### 2.2 Experimental Setups

Flight measurements are conducted within a cubic transparent acrylic observation chamber with an edge length of 30 cm. Captured honeybees are enticed to enter the observation region of the chamber through a transparent conduit. Four high-speed cameras (Phantom VEO 640L, Vision Research, USA) record the honeybees’ takeoff process. The lenses are 105mm fixed-focus lenses (Sigma, Japan). These cameras are strategically positioned above the observation chamber as illustrated in the Figure 1(a) and (b). They operate at a frame rate of 15,000 fps with a resolution of 512×480 pixels. Synchronization of the cameras is achieved through a data transmission link with the built-in trigger system (PTU X, LaVision, Germany) of the controlling computer, ensuring simultaneous capture across all four cameras. To illuminate the honeybees during the experiment, a supplementary lighting system consisting of three LED lights (VISCO, China) with the power of 150 W and the color temperature of 5500 K are employed. Additionally, white diffusion panels are positioned around the observation chamber to facilitate even light distribution and to maintain a white imaging background. This setup is instrumental in enhancing the distinction of various structural elements of the bees in subsequent image processing.

**Figure 1.**
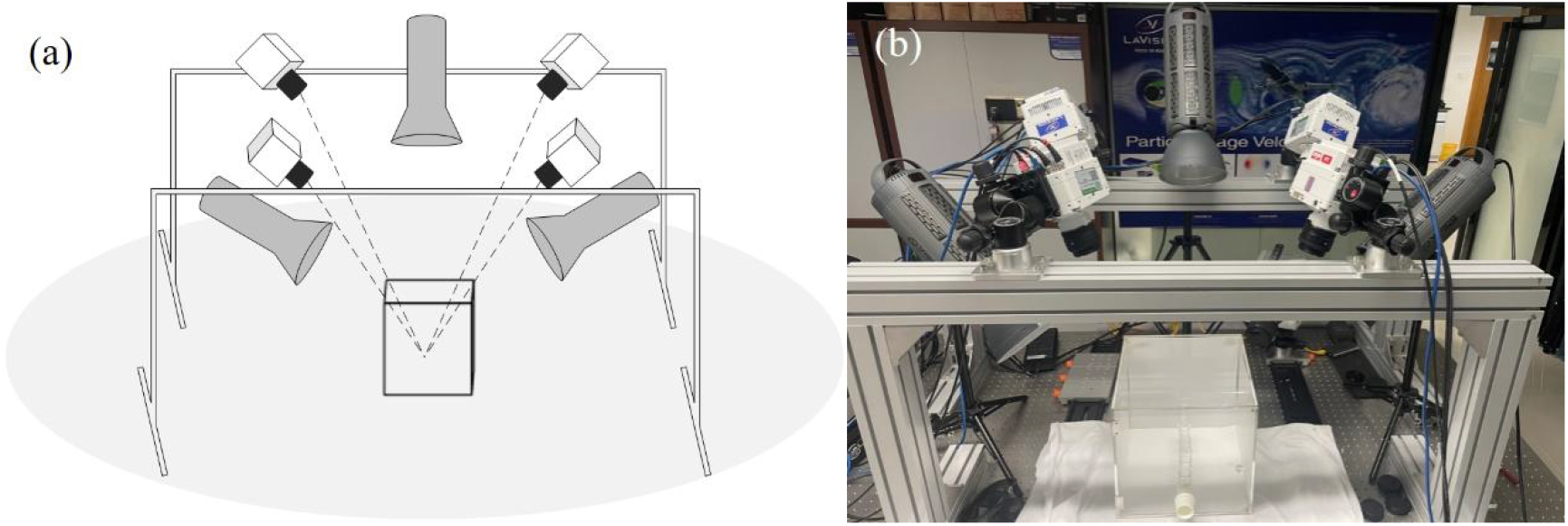
Four high-speed cameras positioned above the observation chamber, with their fields of view converging at the center of the observation chamber. (a) virtual diagram of experimental equipment setup; (b) real setup.

### 2.3 Camera calibration

The calibration method employed in this study is a precise multi-camera calibration method as proposed by Theriault *et al*.^[29]^. A chessboard with 12×9 black-and-white squares was used as the calibration board. By recording the movement of the calibration board within the measurement region, we obtained 300 snapshots of chessboard for each camera. The recorded images then are processed to identify matched image points. The ‘reprojection error’, which refers to the RMS distance between the original and reprojected image points of each calibration point, is 0.063 pixels. The ratio of the standard deviation of wand-length estimates to their mean, is 0.038, which is far below the index value for problems with the calibration. The average uncertainty in the position of each wand tip, estimated from the distance between the two tips is 6.6 × 10^-6^ m. These small errors proved the high accuracy of our calibration.

### 2.4 Wing motion reconstruction

Five points on the wing, either the venation conjunction or the wing tip, are identified as the markers to capture the motion, as shown in Figure 2(a). These markers are on the wing root, leading edge, wingtip, trailing edge, and the juncture between the forewing and hindwing. After recording the wing motion during takeoff of the honeybees, the recorded videos as well as the camera calibration data are loaded into MATLAB software DLTdv8^[30]^ to track these five points frame by frame and extract their three-dimensional coordinates. With the three-dimensional positions of the markers, we simplify the forewing and hindwing to be flat, matching the outline of real honeybee, as shown in Figure 2(b), then align the flat wings with the five markers to reconstruct the flapping of the wings. Note that, as three points determine the plane, markers 1,2,3 are used to align forewing and markers 1,3,4 are used for hindwing. The variation is small when using either point group (1,2,3), point group (1,2,5) or point group (2,3,5) to present forewing. The comparison of pitching angles is shown Figure S1 in Supplementary materials. Overall, ten takeoff flights are processed for analysis.

**Figure 2.**
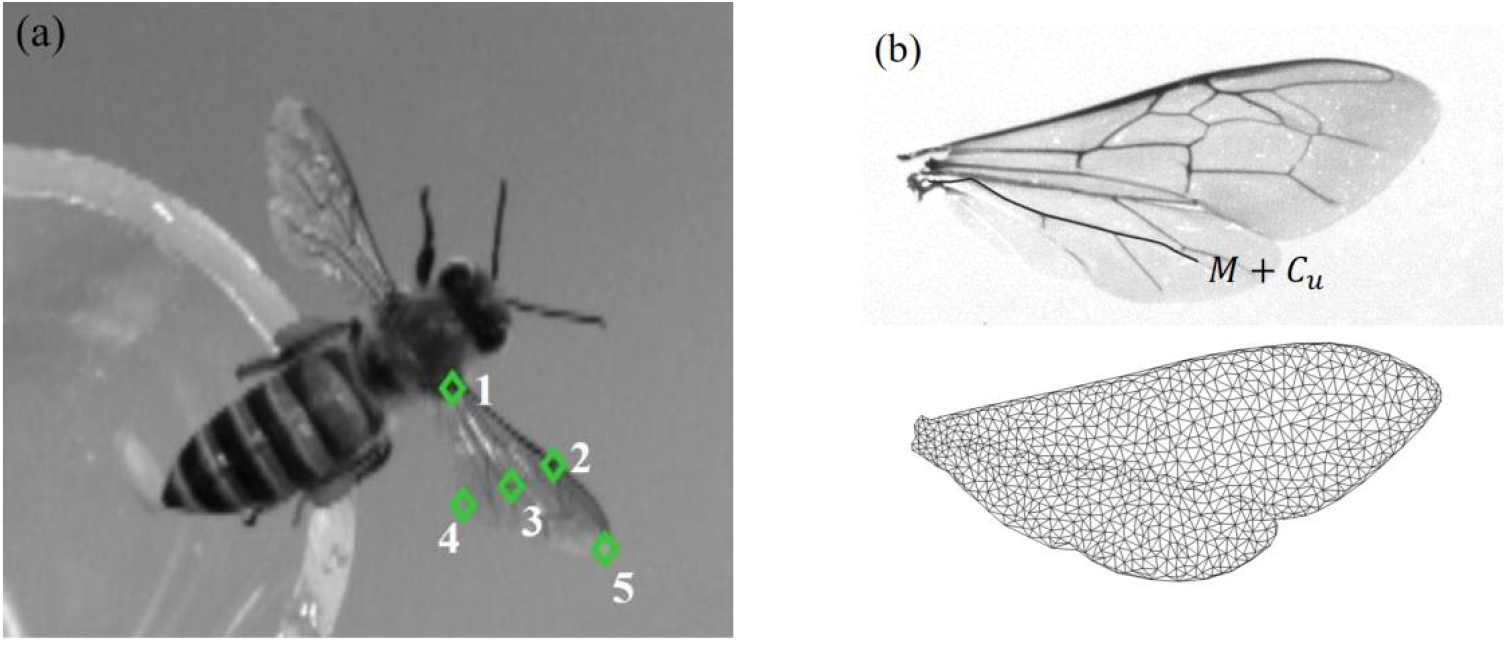
(a): Five tracking points on bee wings, (b): the wing plane model reconstructed from bee wings.

The honeybee’s body is reconstructed using an elliptical cross-section with variable axial lengths. To ensure accurate comparison between the reconstructed wing motion and that of an actual honeybee, we meticulously aligne the reconstructed model with the prototype utilized in its reconstruction. This alignment enables us to observe and analyze both models from an identical perspective, ensuring consistency in our observations. The wing flapping curvature is closely matched, as is shown in Figure 3. Regarding the three metrics mentioned in the previous section for assessing calibration accuracy, the maximal reprojection error of 4 cameras is below 0.1 pixel, the maximal ratio of the standard deviation is below 0.1, and the maximal average uncertainty in the position of each wand tip is below 1.0 × 10^-5^ m. The high-speed videos of one typical takeoff are shown as Video S1-S4 in supplementary materials.

**Figure 3.**
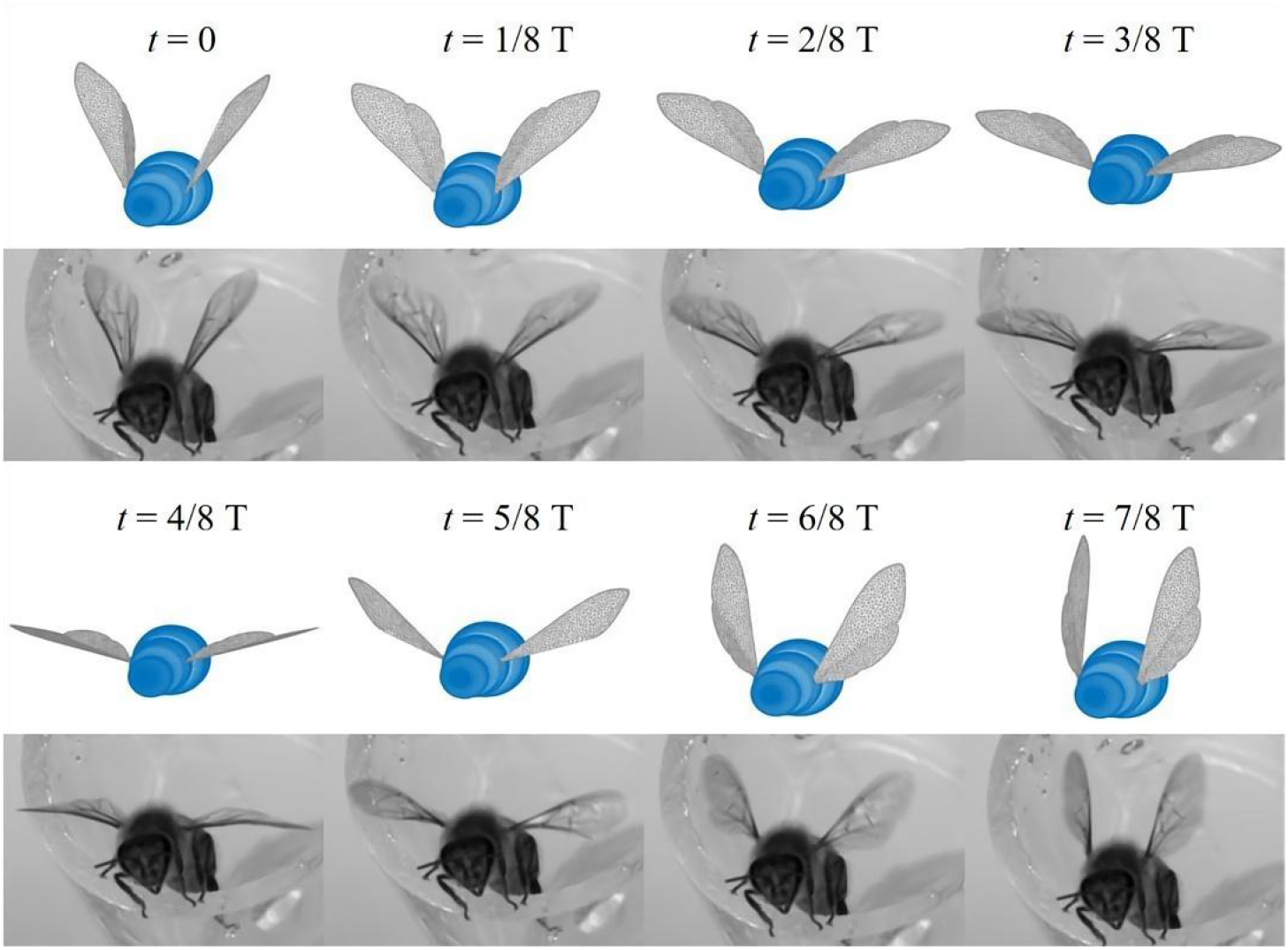
Comparison of reconstruction models and videos at different times in a single cycle

The wing stroke plane is defined by the leading edge of the forewing at its uppermost and lowermost positions. The Euler angles for the forewings, denoted as *Φ*(stroke angle), *θ*(deviation angle), and *Ψ*(pitch angle), are then calculated in the sequence mentioned to determine the position of the forewing, as is shown in Figure 4(a). By determining these Euler angles, we can ascertain the position of the forewing. Subsequently, using the relative positions of the forewing and hindwing, the complete spatial position of the wings can be obtained (Figure 4(b)).

**Figure 4.**
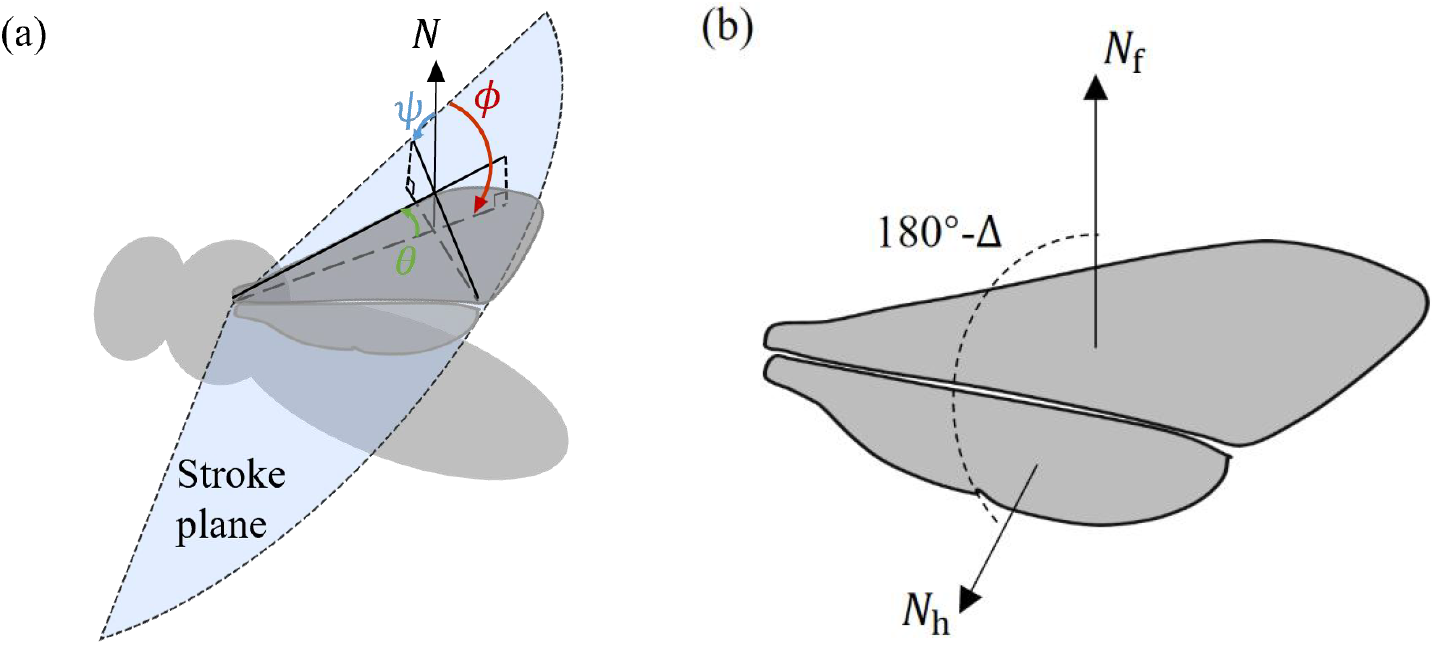
(a) Definition of Euler angles, stroke angle Φ, deviation angle θ and pitching angle Ψ for wing motion, *N* is the normal vector of the stroke plane, (b) angle difference between the forewing and hindwing *N*_f_, is the normal vector of forewing and *N*_h_ is the normal vector of hindwing.

## 3 Results & Discussions

Each takeoff sequence begins from a stationary position and extends to the moment when the bee’s legs leave the surface. When honeybees attach to a surface, the flapping of the wings does not necessarily indicate the initiation of takeoff. In our observations of voluntary takeoff, it is noted that honeybees often flap their wings several times without actually taking off. Statistical analysis of successful takeoffs shows that honeybees typically execute a minimum of 15 wing flap cycles before achieving takeoff.

### 3.1 Division of takeoff to three stages

During takeoff, the variations in the Euler angles exhibit no significant differences among individuals. Figure 5 illustrates the changes in stroke angle, pitch angle, and deviation angle for one typical flight sequence, with the shaded area indicating the downstroke and white area indicating the upstroke. From the figure, it can be observed that variables Φ, *θ*, and *Ψ* exhibit periodic changes over time but with varying amplitude. The wing flapping amplitude increases gradually throughout the takeoff. According to the stroke and deviation angles, the takeoff can be categorized into three stages. The first stage is the initial stage, which usually comprises the first 1-2 beat cycles. In this stage, the amplitude of stroke angle rapidly increases, meanwhile, the deviation angle and pitching angle also increases rapidly. The second stage is the adjusting stage, in which the flapping amplitude and deviation angle continue to increase, transitioning gradually to a larger value. The third stage is the stable stage, which covers the last several beat cycles before detaching the surface. In this stage, the flapping amplitude and deviation angle gradually increased from a large level to the maximum. The angles of other takeoffs shown the similar pattern, as is shown in Figure S2. in supplementary materials. During the last two stages, the pitching amplitude remains stable, as the value reaches 136° in forth cycle and variation is within 15° during the remaining cycles. Throughout the takeoff, the angle between the stroke plane and the body is approximately 110°.

**Figure 5.**
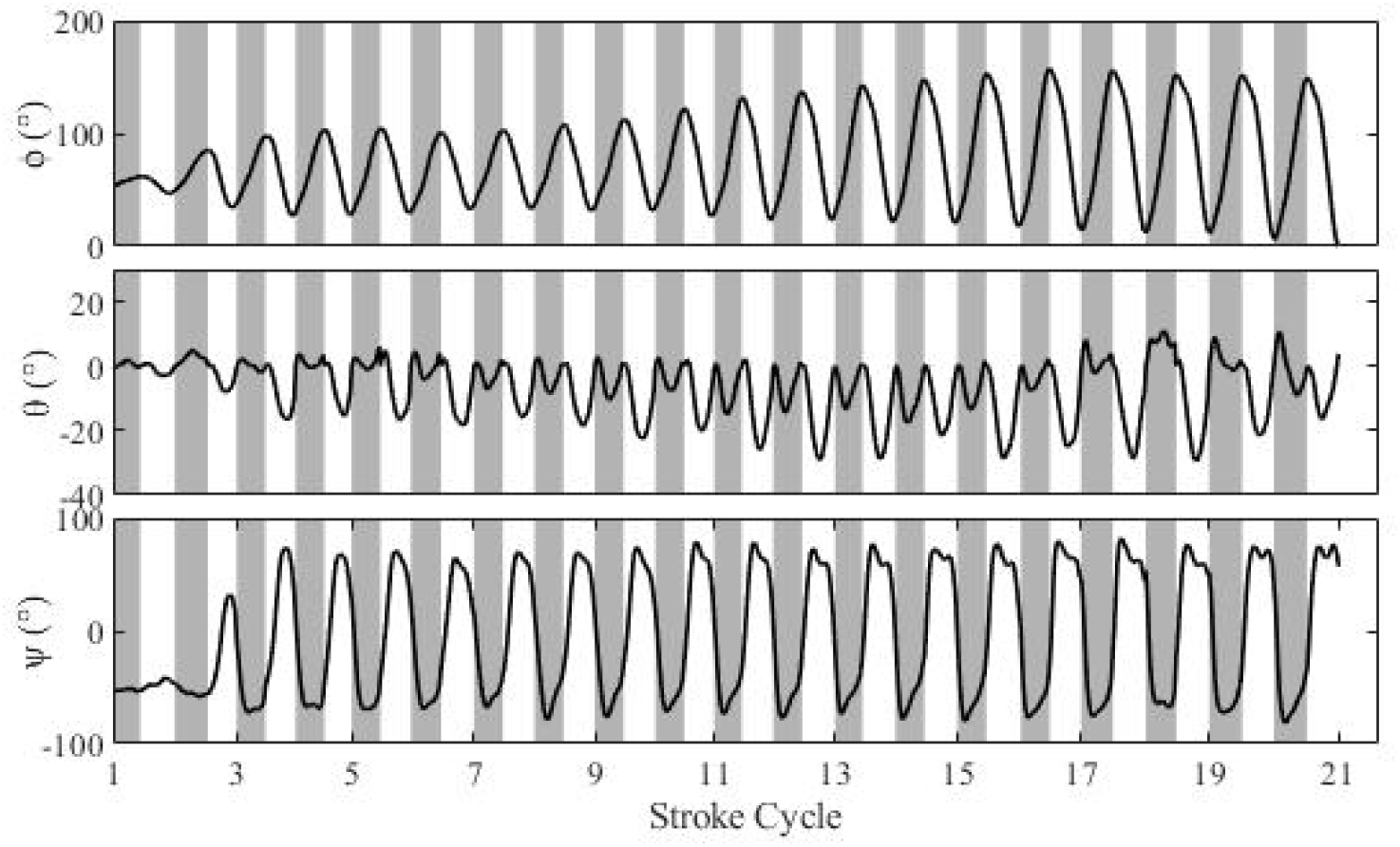
The changes of stroke angle (ϕ), deviation angle (θ) and pitch angle (ψ) over time, with shaded areas indicating the downstroke

The wingtip trajectories of honeybees in different stages are shown in Figure 6. The angle between the stroke plane and the body does not exhibit significant changes throughout the takeoff process. In the initial stage, the wingtip motion trajectories resemble ellipses, with the wingtip trajectory being above the stroke plane during downstroke and below it during upstroke. In the adjusting stage, where both deviation angles become negative, the wingtip trajectories shift predominantly below the stroke plane. In the stable stage, the downstroke trajectories intersect with the upstroke trajectories, forming a figure-of-eight shape trajectory. The deviation angles continue to decrease, causing the wingtip trajectories to deviate further from the stroke plane. In the simulation conducted by Lei *et al*. of models with different deviation angles, it is found that the figure-eight flapping pattern produces the highest lift coefficient, while the zero-deviation flapping pattern achieved the highest lift-to-power ratio^[31]^. Luo *et al*. Points out that deviation typically leads to an increase in energy consumption during insect flight, indicating that insects tend to maintain flapping motion during flight^[32]^.

**Figure 6.**
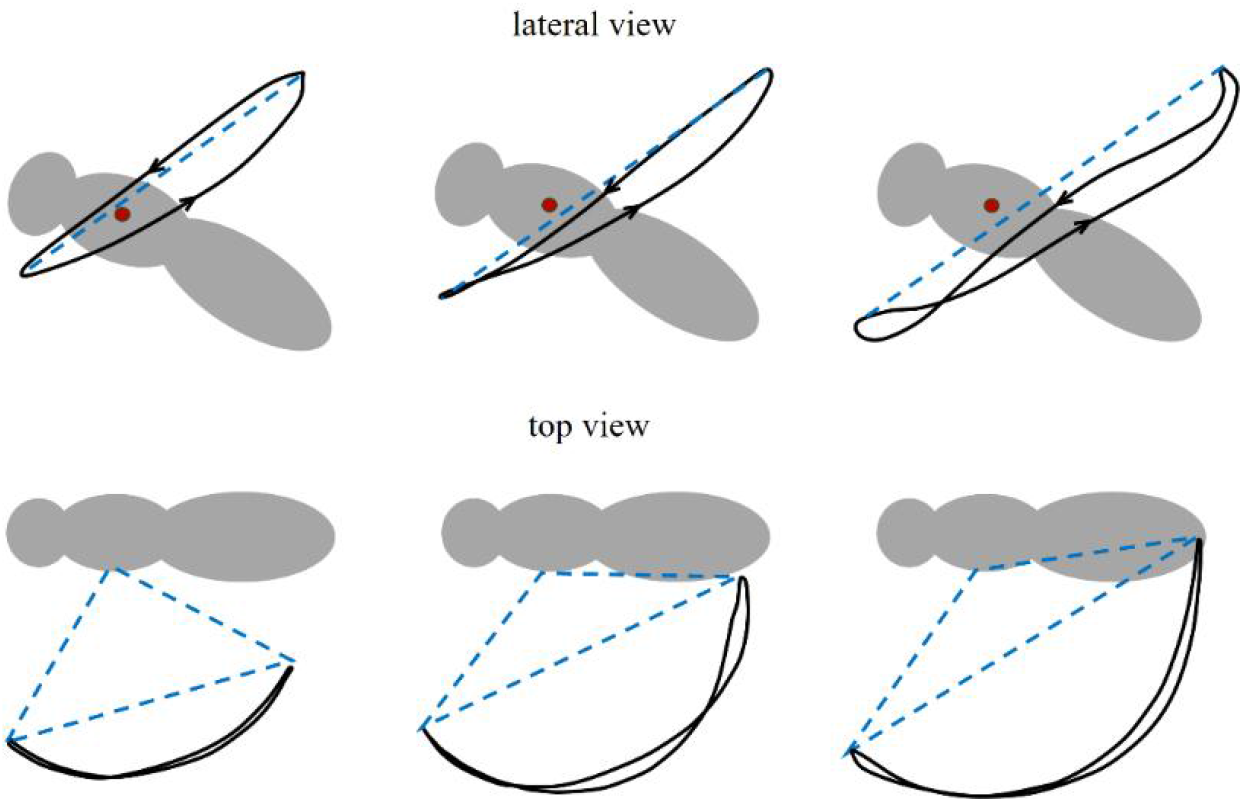
Wingtip trajectories in three stages of deviation angle, the blue dashed line represents stroke plane and the red dot indicates the position of the wing root.

### 3.2 Wingbeat frequency, flapping amplitude, and wingtip velocity

During takeoff sequences, the number of cycles prior to detachment from a surface varies between, and within, individuals. For direct comparison, we analyse the last 10 cycles prior to last contact. Figure 7 (a-c) shows the wingbeat frequency (one over stroke period of each cycle), flapping amplitude and cyclic average wingtip speed, and Figure 7(d) shows correlations between these parameters. In the measurement, nine of the takeoff flights showed that the wingbeat frequency exhibit a relatively small difference during the last 10 cycles within an individual takeoff sequence, with an average increase of 20 Hz. In one exceptional flight, the frequency increased from 120 Hz to 160 Hz, and its detaching frequency is still smaller than the others. At the last cycle, the wingbeat frequency ranges from 180 Hz to 260 Hz, with the average 220 Hz. In comparison, the stroke amplitude increases during the last ten cycles, with the detaching amplitude ranging from 110° to 140°. Generally, the last several cycles maintain the similar amplitude, indicating a stable stage, while the gradually increasing cycles indicate a adjusting stage.

**Figure 7.**
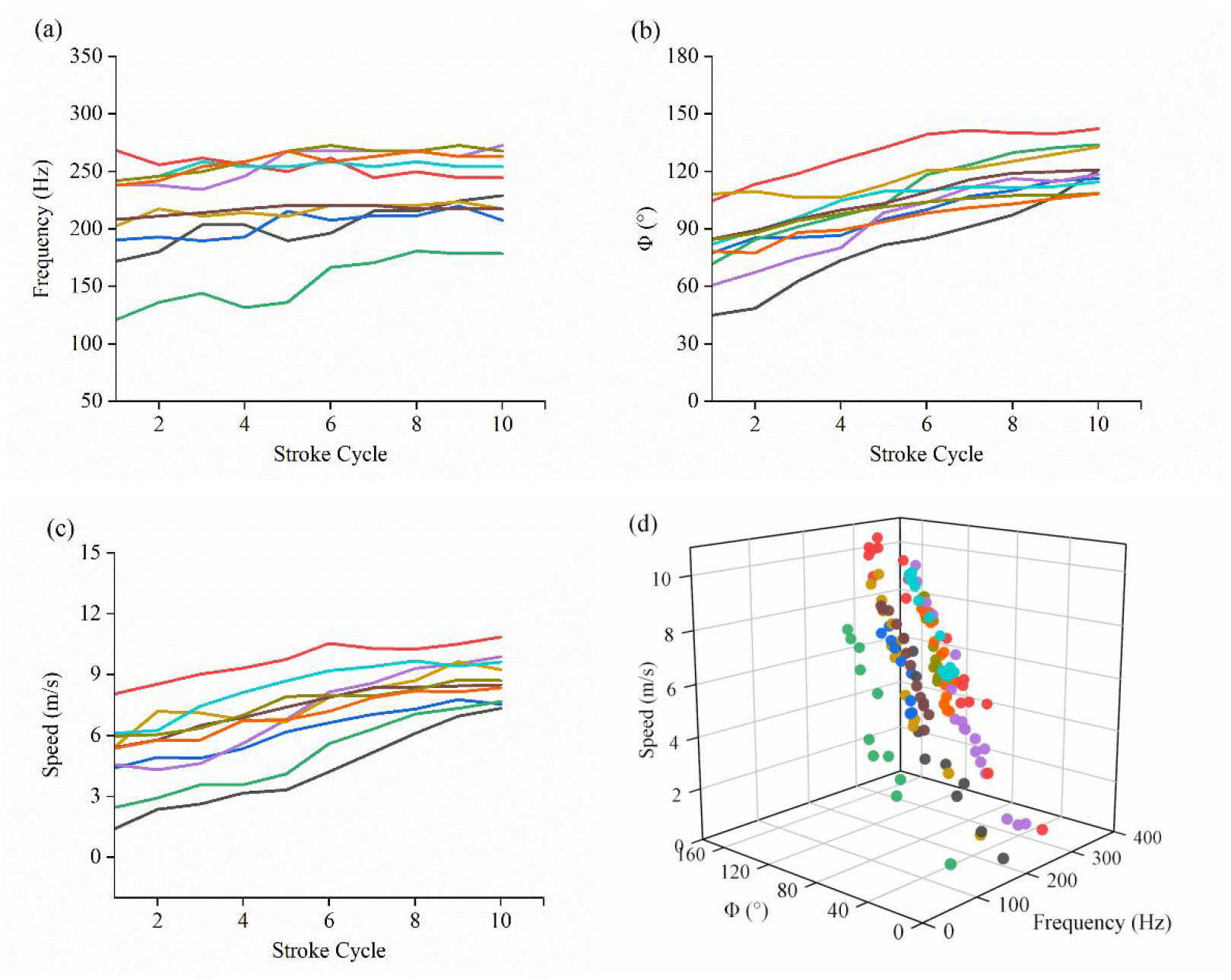
(a) The variation of wingbeat frequency (one over stroke period in each cycle), (b) flapping amplitude, and (c) wingtip cycle-average velocity over the wingbeat cycles, (d) the relationships between wingtip average velocity, wingbeat frequency, and flapping amplitude.

The wingtip speed is calculated with the 3D tracking position, and the peak instantaneous wingtip velocity upon detaching the surface reaches approximately 14.7± 1.7 m/s for all takeoffs. The cyclic wingtip speed during the last ten cycles shows the increase in magnitude with the cycle. The cycle-average speed ranges from 7.3 m/s to 10.8 m/s with the mean value of 8.7 m/s. The wingtip speed is related to both frequency and stroke amplitude with the equation U_t_ = 2 Φ*f*R, where Φ represents stroke amplitude, *f* stands for wingbeat frequency, and R is the wing length. Figure 7(d) displays the correlation between wingbeat frequency, stroke amplitude, and wingtip speed for all cycles in the ten flight sequences. The Pearson correlation coefficient between frequency and speed is 0.5, while between amplitude and speed is 0.9, indicating that changes in wingtip average speed are primarily influenced by variations in flapping amplitude.

### 3.3 The camber between forewing and hindwing

During flight, the honeybee’s forewing and hindwing are coupled by means of the forewing posterior rolled margin and hindwing hamuli^[27]^. Due to this structural arrangement, they can be thought of as functionally two-winged insects, and the aerodynamics likely to be similar to those of dipterous insects. However, the hinge-like connection between the forewing and hindwing reduces the effective stiffness in the chordwise direction, resulting in a large chordwise deformation on each stroke. In this study, the variation in the angle between forewing and hindwing, Δ, is analyzed. Figure 8(a) illustrates the cyclic average of Δ over the last five cycles, with minimal difference between cycles. Overall, Δ undergoes positive values at downstroke and negative values at upstroke, meaning the combined wings make positive camber during both the downstroke and upstroke portions of the stroke cycle. The downstroke and upstroke of the wingbeat cycle exhibit a similar variation pattern, with the angle of deflection Δ initially increasing to its maximum value of 54° and -56° at t/T = 0.34 and 0.6 for downstroke and upstroke, respectively Subsequently, the angle of deflection decreases and oscillates around 21° in the downstroke and -37° in the upstroke before reversing sign at stroke reversal. This pattern is comparable to that observed in dipterous hoverflies, where the camber also reaches a peak early in downstroke (t/T = 0.07) and upstroke (t/T = 0.54), then decreases and rebounds to a second peak, following by a sharp transition ^[23].^ This behavior is believed to be driven by passive dynamics influenced by wing inertia^[33]^. In the study of tethered honeybee flight, Ma et al.’s raw data also indicated this pattern, which was previously overlooked due to the use of a first-order sinusoidal fitting applied over the entire cycle^[27]^. Our analysis of honeybee flight videos suggests that this phenomenon results from rapid wing flipping, causing an inertial lag in the hindwings (refer to Figure S3 and Video 5 in supplementary materials). The detailed mechanism behind this observation is not within the scope of this paper and will be the subject of future investigation.

**Figure 8.**
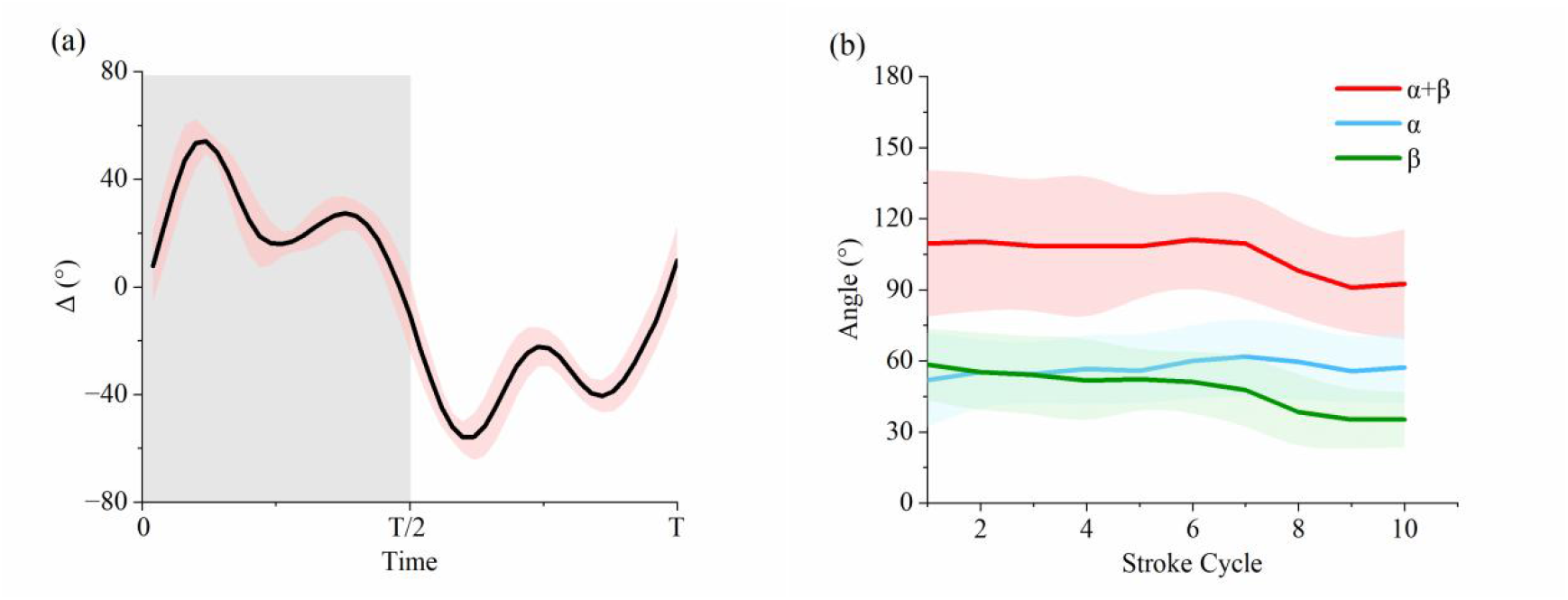
(a) The temporal variations in the angle between forewing and hindwing. (b) average value of maximum convex angle (α), maximum concave angle (β) and α+β with their error band in all flights.

We define the maximum convex angle at downstroke as α and designate the largest concave angle amplitude at upstroke as β. Figure 8(b) illustrates the variation of mean values of 10 takeoffs and their error bands for α and β in last 10 cycles. Throughout the entire takeoff process, the maximum convex angle in each cycle progressively increases, while the concave angle amplitude decreases. α + β can be used to represent the amplitude of Δ changes. The large error margin of α + β indicates that significant variations exist in the Δ amplitudes between individuals or takeoffs. From Figure 8(b), it can be seen that the amplitudes of Δ, or α + β, remain unchanged firstly and decrease subsequently prior before detaching the surface.

### 3.4 Camber effect on aerodynamics

The effect of the camber on the aerodynamics of insect wings is disputable. Prior research focusing on various species, including locusts, Drosophila, bumblebees, and hummingbirds, has suggested that camber increases maximum lift coefficients and improves the lift-to-drag ratio^[34-37]^. However, other studies examining insect wings in unsteady flow conditions have found that the aerodynamic benefits conferred by camber are minimal^[38, 39]^. Contrasting these findings, a numerical analysis of cicada wing flapping demonstrated that pre-existing camber significantly enhances thrust during the downstroke^[40]^ and effectively reduces downward force during the upstroke while the flow is also unsteady^[41, 42]^. Our study focuses on the hinged wings of honeybees, which demonstrate positive cambers during both downstroke and upstroke on different stage of takeoff. As shown in Figure 9(a), a chordwise wing profile displays the positive camber throughout the cycles in both growth and stable stages.The camber enhances effective angle of attacks, implying some aerodynamic advantages.

**Figure 9.**
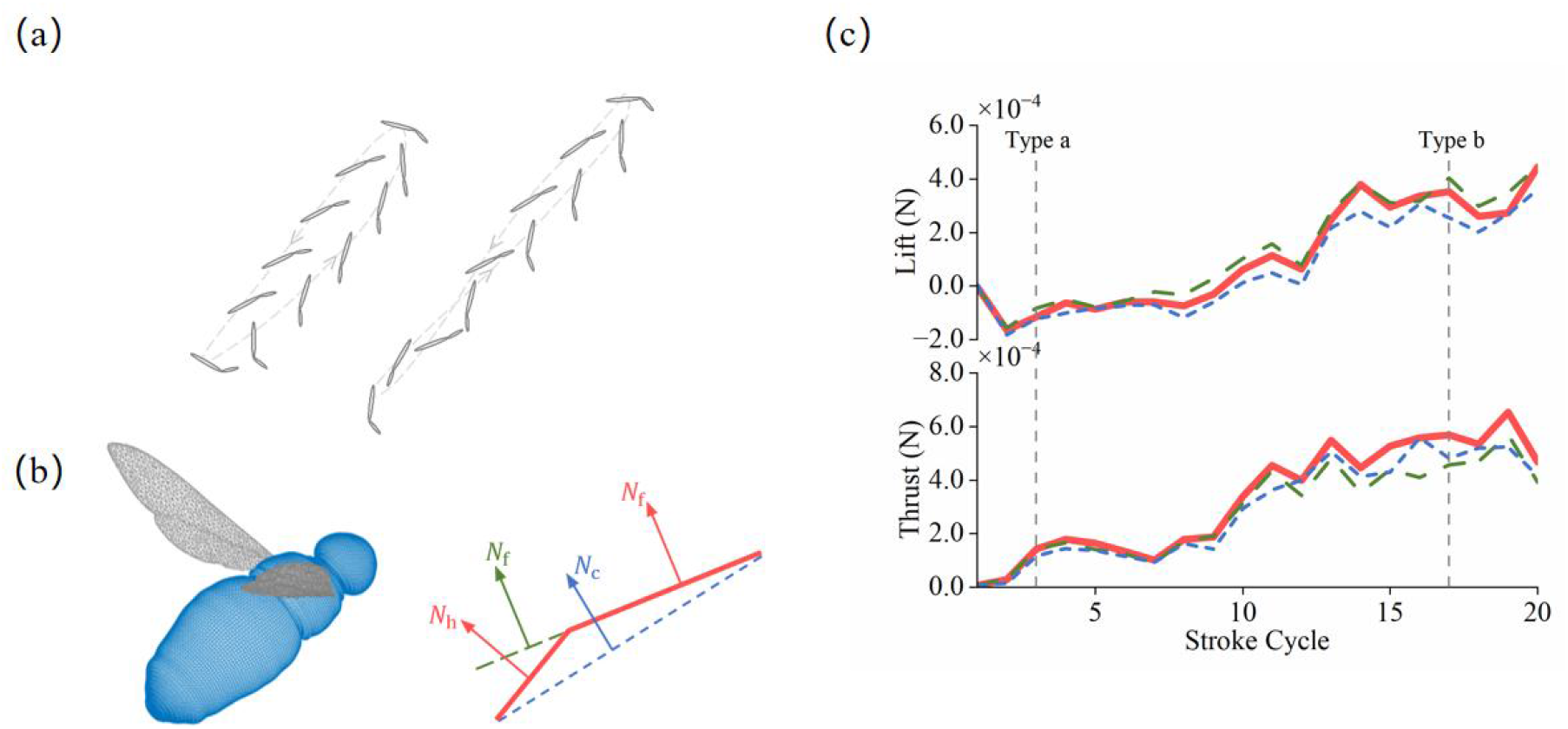
(a) Simplified wing chordwise profile in two types of wingtip trajectories on different stage of takeoff. (b) the illustration of three wing models, including a cambered wing from high-speed videography, and two flat wings.(c) The cycle-averaged lift and thrust generated by one wing throughout the takeoff.

To determine the aerodynamic effects of positive wing camber, we employ the immersed-boundary method to simulate the forces acting on both cambered (right wing) and two corresponding flat wings: one mirroring the normal vector **N**_**f**_ of the forewing and the other combining the normal vectors **N**_**f**_ and **N**_**c**_ of both the forewing and hindwing, as depicted in Figure 9(b). The results reveal that the cambered wing does not significantly contribute to lift during takeoff; however, it does facilitate a notable enhancement in thrust, as shown in Figure 9(c) and the temporal variation and flow shown in Figure S4 in Supplementary materials. The average lift and thrust values during the adjusting and stable stages are listed in Table 2. We observe that the lift generated by the cambered wing is comparable to that of the flat wings in both stages. In terms of thrust, the cambered wing during the adjusting stage produces values 9.0% and 12.7% higher than those of the respective flat wings, and 18% and 11% greater during the stable stage. This finding highlights the critical role of forward motion in certain types of takeoffs^[40]^.

**Table 2.** The average lift and thrust generation of reconstructed cambered wing and two flat wings on adjusting and stable stages.

| Wing models | Lift ($\times 10^{-4}$ N) | | Thrust ($\times 10^{-4}$ N) | |
| --- | --- | --- | --- | --- |
|  | Adjusting | Stable | Adjusting | Stable |
| Cambered wing | -0.13 | 3.35 | 2.20 | 5.36 |
| Flat wing $N_f$ | 0.13 | 3.60 | 2.00 | 4.42 |
| Flat wing $N_c$ | -0.40 | 1.70 | 1.92 | 4.76 |

### 3.5 Actuation of hindwing pitching

Flat flapping wings constructed from isotropic materials exhibit a tendency for the trailing edge to move backward more significantly, resulting in a reverse camber due to aerodynamic and inertial effects^[43, 44]^. This passive deformation behavior, influenced by factors such as wing stiffness and mass ratio, allows for the periodic storage and release of energy during flapping, thereby enhancing aerodynamic efficiency^[45]^. Furthermore, as the wings undergo periodic acceleration and deceleration, both inertial and aerodynamic forces contribute to the pitching motion of the wings^[46]^, a phenomenon observed in dragonflies, hawkmoths, and fruit flies ^[47, 48]^. In this process, the wing root acts as a torsional damping mechanism^[33, 48]^. Despite similarities in flapping motions between honeybees and other insects like the fruit fly, the specific mechanisms of honeybee wing pitching remain elusive. This complexity is partly due to the connection of the hindwing to the forewing via the hamuli structure, which complicates dynamics due to the coupling of the two wings. The actuation of the hindwing in honeybees is particularly intriguing. While anatomical studies reveal that honeybee hindwings are attached to direct muscles at their base, enabling actuation^[49]^, it is still unclear whether these muscles are used to drive the pitching motion.

To address this question, we combined inertial and aerodynamic forces for a qualitative analysis of a cycle in the stable stage of takeoff. Figure 10(a) illustrates the temporal variation of tip velocity (*v*_t_), tip tangential acceleration (a_t_), and the angular acceleration of the forewing along its leading edge during the stable stage. If we establish a forewing-fixed coordinate system (oxyz), it is important to note that this system is non-inertial. During pronation and supination, the angular acceleration is substantial, potentially generating a significant inertial force (**F**_**I**_) that contributes to forming the positive camber, as shown in Figure 10(b,i). This mechanism, stemming from the inertial effect, likely aids in establishing positive camber at the start of each half-stroke, where angular acceleration is pronounced, without requiring any actuation at the base of the hindwing. As the wing progresses to the middle of the downstroke/upstroke, the pitching acceleration rapidly decreases. Here, both the aerodynamic force (**F**_**a**_) and inertial force (**F**_**I**_) contribute to an anti-clockwise rotation of the hindwing, due to the large velocity (*v*_t_) and some tangential acceleration (a_t_). This phase challenges the maintenance of positive camber, as depicted in Figure 10(b, ii), necessitating actuation at the root of the hindwing. This can be achieved either by applying a pitching torque (**M**) on the leading edge of the hindwing or exerting a translational force (**F**) along the *M+Cu* vein of the hindwing (shown in Figure 2(b)). Either hypothesis could explain the sustained positive wing flexion during flapping, and it’s plausible that honeybees might use these methods independently or in combination.Following the middle stroke, the aerodynamic and inertial forces can induce pitching torque in opposite directions, as shown in Figure 10(b,iii). However, the aerodynamic effect weakens due to changes in *v*_t_, while the inertial effect strengthens in the opposite direction, leading to a gradual transition from active to passive pitching of the hindwing. This qualitative analysis suggests that the hindwing should be actively actuated for pitching before and around the middle of the half-stroke, whereas passive pitching occurs at other times. Future research will more precisely explore the role of thoracic muscles in this process.

**Figure 10.**
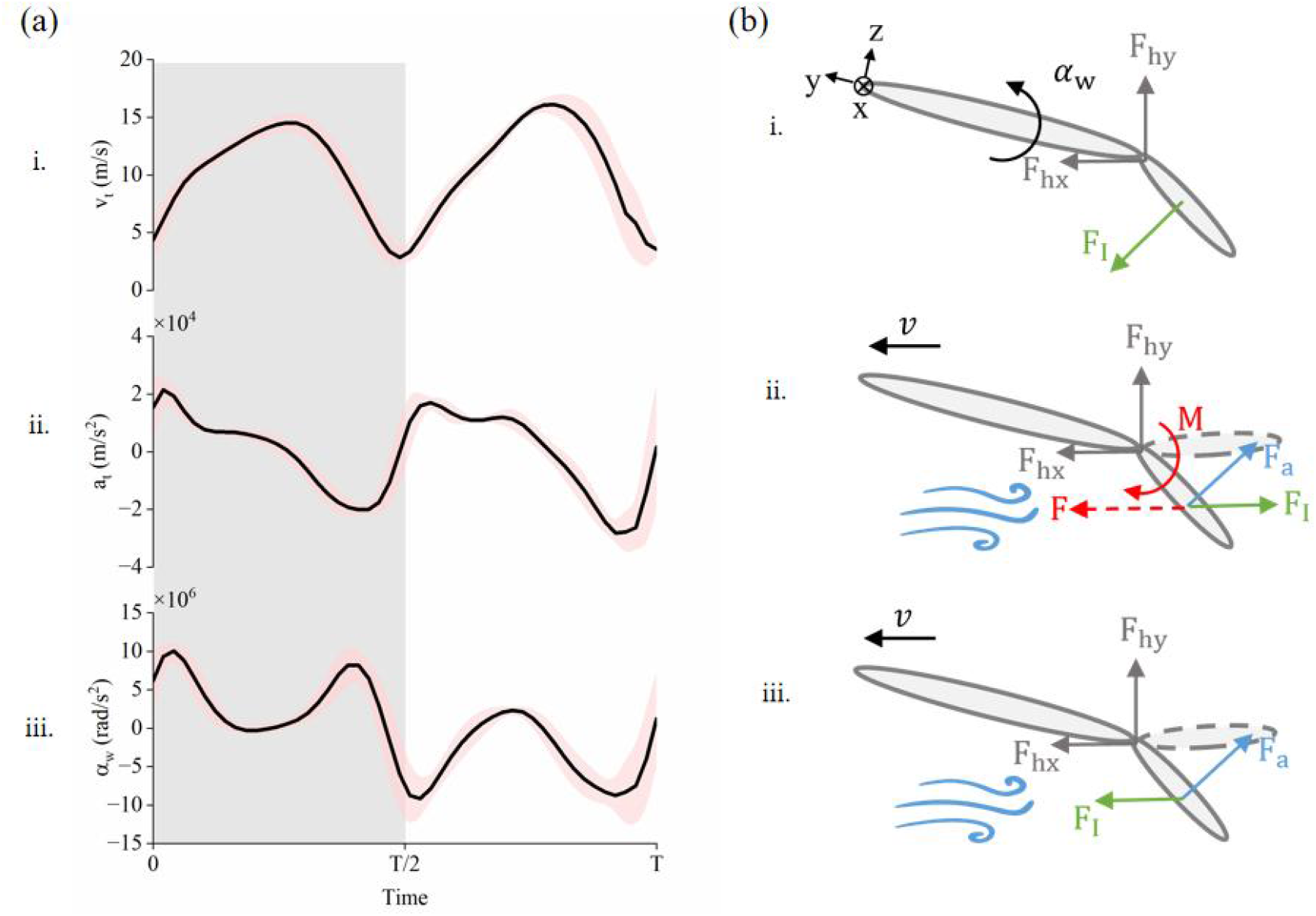
(a) The cycle-averaged temporal variation of wing tip velocity *v*_t_ (i), tip tangential acceleration a_t_ (ii) and the angular acceleration of the forewing along its leading edge (iii) on stable stage; (b) typical force configuration on the hindwing and its possible actuation. (i) during pronation and suppination; (b) prior and at middle downstroke/upstroke,(iii) posterior to middle downstroke/upstroke. F_hx_ and F_hy_ are the forces exerted by hamuli on the hindwings, **F**_**a**_ is the aerodynamic force, **F**_**I**_ is the inertial force, **M** is the pitching torque along leading edge and **F** is the translational force exerted on hindwings by the thoracic muscles.

## 4 Conclusions

This research utilize four high-speed cameras to meticulously capture the takeoff process of several honeybees, leading to the development of a detailed motion model for honeybee takeoff. Kinematic analysis identifies three distinct stages of takeoff, characterized by variations in stroke and deviation angles. An examination of the final ten wingbeat cycles before takeoff reveals a constant frequency, while stroke amplitude and wingtip velocity progressively increased. Notably, a positive aerodynamic camber is observed during both downstroke and upstroke, accompanied by a midpoint recoil in each cycle, although the mechanics behind this remain uncertain. Our numerical findings suggest that this positive camber predominantly contributes to increased thrust rather than lift during takeoff, underscoring its importance in facilitating forward flight. Moreover, it appears that muscular action is required to initiate and maintain wing motion at the beginning and middle of each downstroke and upstroke, with the remainder of the motion being passive. This study illuminates the potential of employing hinged-wing mechanisms in the design of flapping wing micro air vehicles, drawing inspiration from the wing motion of honeybees.

## Acknowledgement

This work is supported by the Natural Science Foundation of Guangdong Province (No. 2023A1515011506) and National Natural Science Foundation of China (No. 12302296) to JS, the Guangdong Provincial Key Laboratory of Intelligent Disaster Prevention and Emergency Technologies for Urban Lifeline Engineering (No.2022B1212010016) to LY and JS, and the National Natural Science Foundation of China (No. 52005104) to CW.

