## Supplementary Figure S1-S4 for "The kinematics and aerodynamics of hinged wing of honeybees during takeoff": Supplementary materials.docx

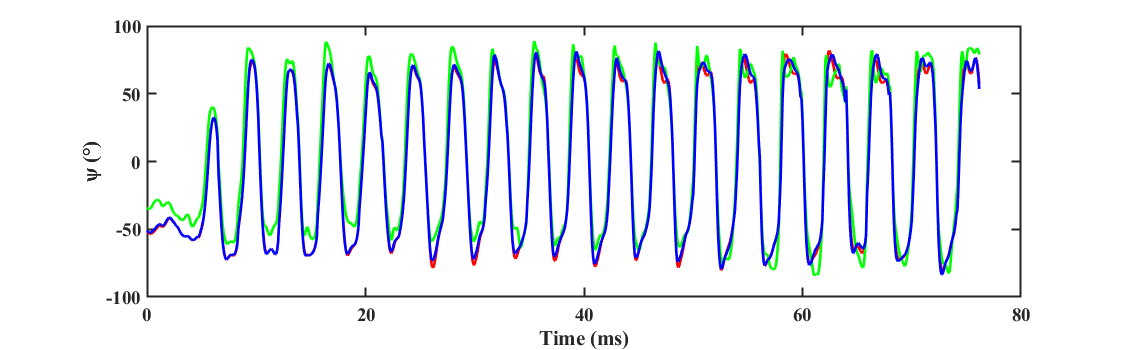

**Figure S1**. The comparison of pitching angle using point group (1,2,3) (red line), point group (1,2,5)(green line), and point group (2,3,5)(blue line). The corresponding magnitudes are ?, and ?? and ??.

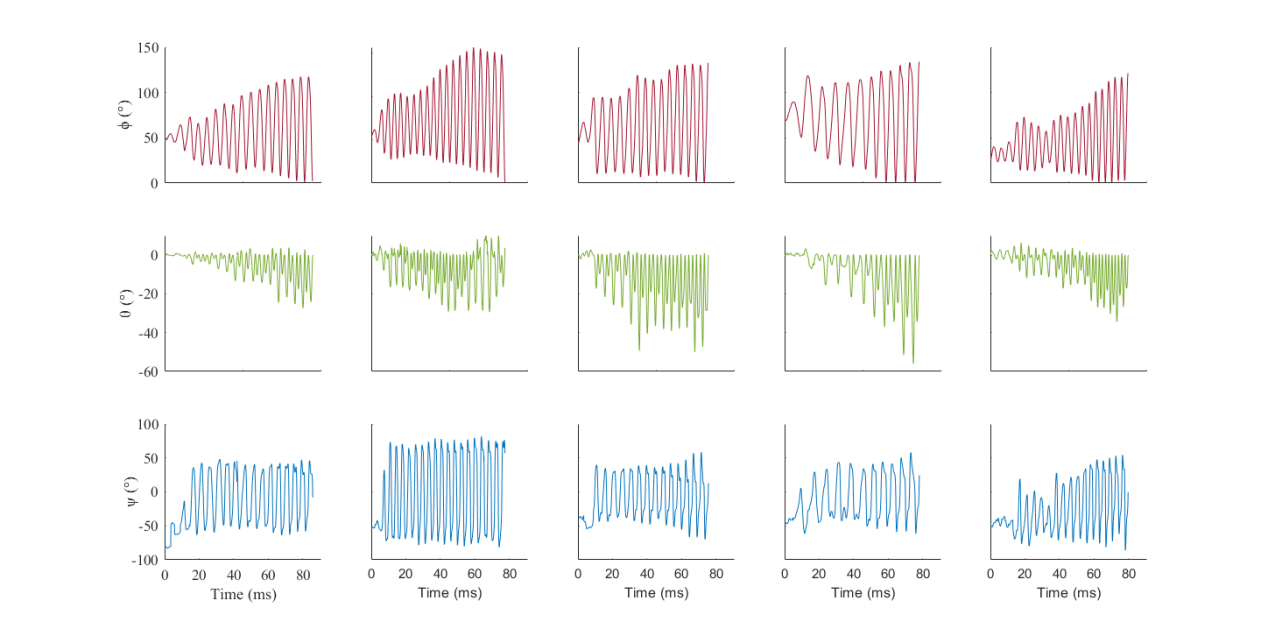

**Figure S2.** The changes of Euler angles stroke angle (ϕ), deviation angle (θ) and pitch angle (ψ) over time of five flights. (Envelop of three angles )

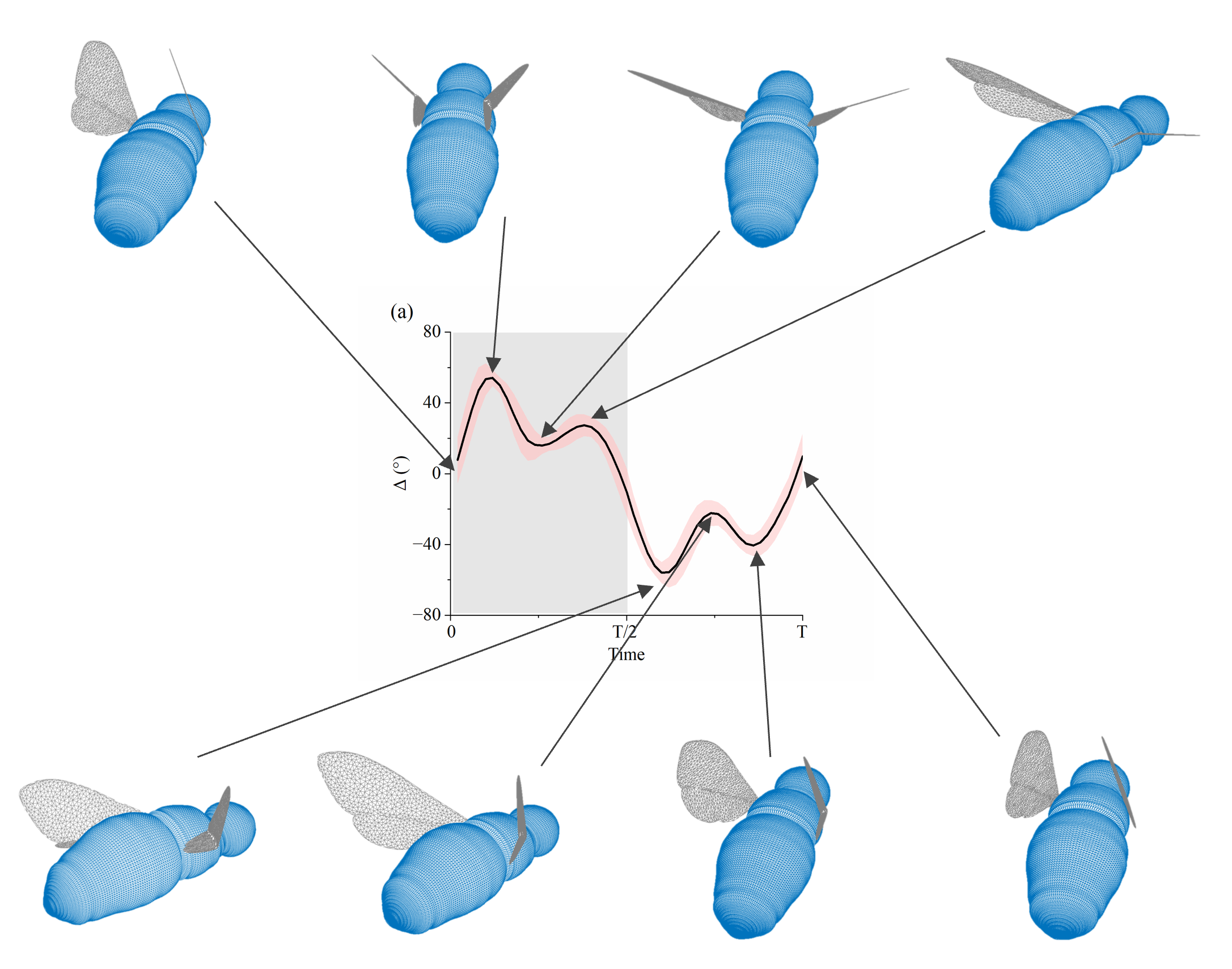

**Figure S3.** The illustration of hindwing recoil with 8 snapshots during both downstroke and upstroke.

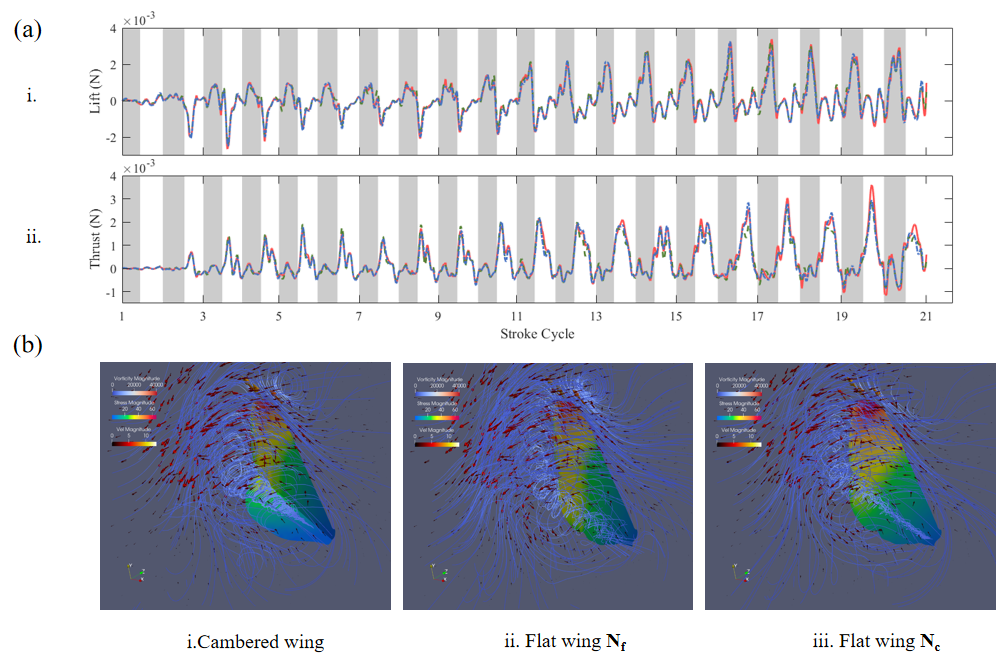

**Figure S4.** (a) The temporal variation of thrust and lift during the takeoff of a honeybee with three wing models, the red line indicates the cambered wing from, and the green line indicates the flat wing with same pitching as the forewing, and the blue line indicate the flat wing with combined pitching; (b) the flow characters of three wing models.
