## Supplementary figures and images for "The kinematics and aerodynamics of hinged wing of honeybees during takeoff"

### Supplementary Video S5

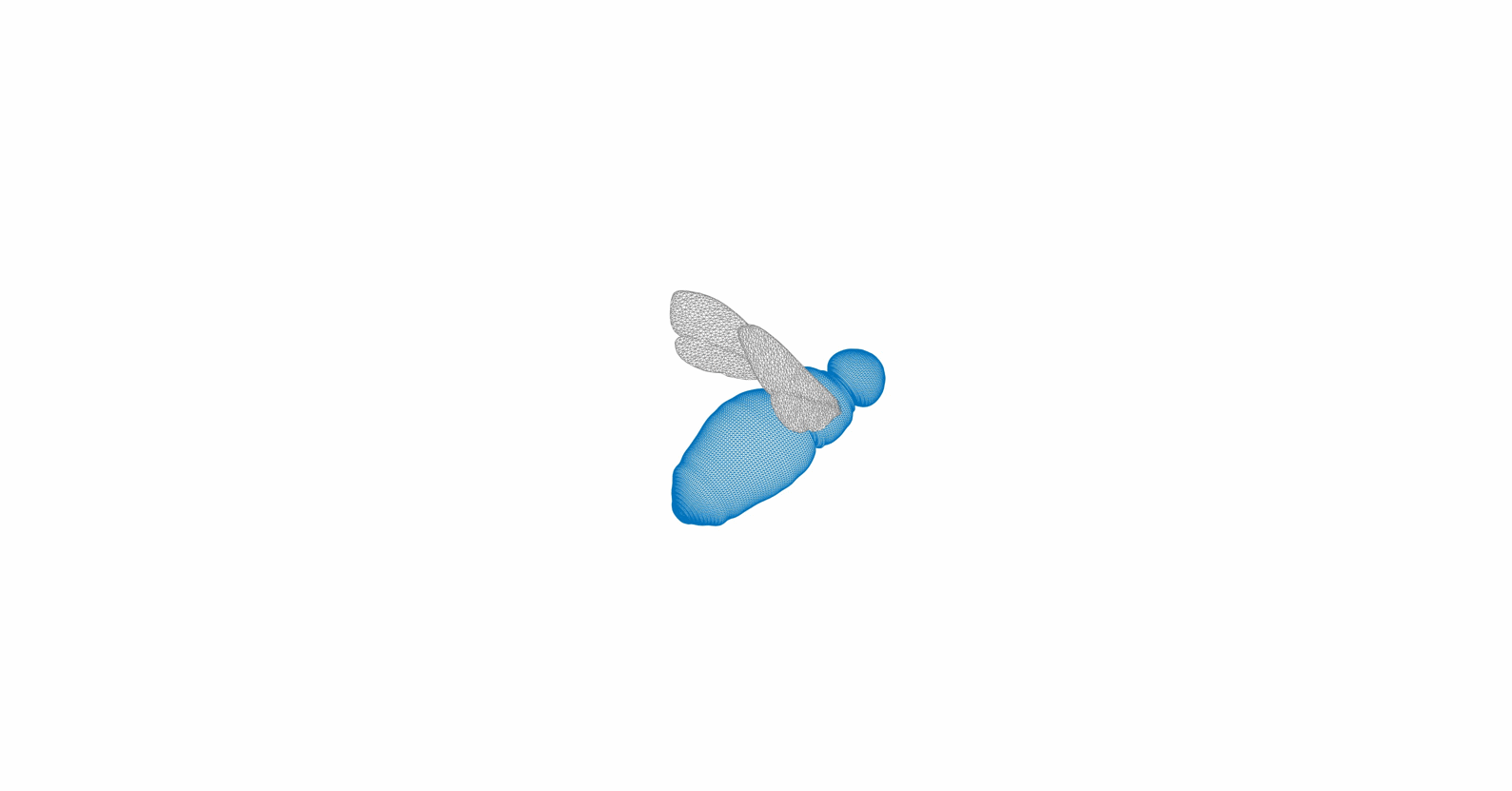
